# A Witches’ Broom Phytoplasma effector induces stunting by stabilizing a bHLH transcription factor in plants

**DOI:** 10.1101/2024.12.24.629987

**Authors:** Shuang Yang, Amelia Lovelace, Yi Yuan, Haizhen Nie, Weikai Chen, Yi Gao, Wenhao Bo, Dawn H Nagel, Xiaoming Pang, Wenbo Ma

**Affiliations:** State Key Laboratory of Tree Genetics and Breeding, National Engineering Research Center of Tree Breeding and Ecological Restoration, Key Laboratory of Genetics and Breeding in Forest Trees and Ornamental Plants, Ministry of Education, College of Biological Sciences and Biotechnology, Beijing Forestry University, Beijing, 100083, China; The Sainsbury Laboratory, Norwich Research Park, Norwich, NR4 7UH, UK; Department of Botany and Plant Sciences, University of California Riverside, Riverside, CA 92521, USA

## Abstract

- Phytoplasmas are specialized phloem-limited bacteria that cause diseases on various crops resulting in significant agricultural losses. This research focuses on the jujube Witches’ Broom (JWB) phytoplasma and investigates the host-manipulating activity of the effector SJP39.
- We found that SJP39 directly interacts with the plant transcription factor bHLH87 in the nuclei. SJP39 stabilizes the bHLH87 homologs in *A. thaliana* and jujube, leading to growth defects in the plants.
- Transcriptomic analysis indicates that SJP39 affects the gibberellin (GA) pathway in jujube. We further demonstrate that ZjbHLH87 regulates GA signalling as a negative regulator and SJP39 enhances this regulation.
- The research offers important insights into the pathogenesis of JWB disease and identified SJP39 as a virulence factor that can contribute to the growth defects caused by JWB phytoplasma infection. These findings open new opportunities to manage JWB and other phytoplasma diseases.

## Introduction

Jujube (*Ziziphus jujuba* Mill.) is one of the longest-cultivated perennial fruit crops originated in China (Li et al., 2024) with significant impacts in culture and economy. The jujube industry is currently facing an unprecedented threat due to the Jujube Witches’ Broom (JWB) disease, which affects main jujube cultivation areas in China (Guo et al., 2023). Diseased jujube trees exhibit a spectrum of symptoms including witches’ broom, phyllody, leaf yellowing, and stunting trees (Bertaccini, 2022; Guo et al., 2023). Infected trees typically succumb to the disease within 2-3 years (Zhou et al., 2021). Economic losses caused by JWB are not only due to reduced yield, fruit quality, and tree longevity, but also substantial costs for disease control and management measures (Wang et al., 2018a; Yang et al., 2023). Thus, a deep understanding of the virulence mechanisms underlying JWB development is crucial for developing effective disease control strategies and to safeguard the global jujube industry.

JWB is caused by phytoplasma infection. Phytoplasmas are obligate bacterial pathogens that colonize phloem sieve elements of infected plants through insect transmission (Ye et al., 2017). Phytoplasmas undergo a life cycle that alternates between plant hosts and specific insects (Wei et al., 2004). Insects become infected by acquiring phytoplasmas from the phloem of diseased plants. Once the bacteria invade the insect’s salivary glands, they can be transferred to the phloem of healthy plants during feeding (Namba, 2019). These bacteria lack a cell wall and possess a highly reduced genome, reflecting their lifestyle as obligate pathogens (Oshima et al., 2013; Guo et al., 2023). The phytoplasma responsible for JWB belongs to the 16SrV-B sub-group (Xue et al., 2023). The primary insect vector for JWB phytoplasma transmission is the leafhopper *Hishimonus sellatus* (Jung et al., 2003). In addition, JWB can also spread through grafting practices (Lee et al., 2012; Ye et al., 2017). Currently, phytoplasmas axenic culture of phytoplasmas is difficult, and few chemical or biological control methods exist for JWB disease management. Previous studies have demonstrated that techniques such as shoot tip culture to produce pathogen-free propagations (Wang et al., 2009; Namba, 2019) and injecting tetracycline derivatives into infected stems or seeds can reduce phytoplasma infection in plant explants (Askari et al., 2011; Bertaccini, 2022). However, these management measures are costly, laborious, and time-consuming. While there is an urgent need to develop novel approaches to accomplish sustainable resistance to JWB in jujube, achieving this goal requires a comprehensive understanding of the disease and the phytoplasma pathogen.

During the infection process, phytoplasma secretes a variety of effector proteins via the Sec-dependent pathway (Gao et al., 2023). These effectors, which are crucial for the phytoplasma colonization, insect transmission, and disease symptom development.

To date, approximately 20 phytoplasma effectors have been experimentally studied, although the number of predicted effector proteins is much larger (Carreón et al., 2023). The best-studied effectors are encoded in the strain Aster yellows witches’ broom (AY-WB; or *Ca* Phytoplasma asteris), which infects the model plant *Arabidopsis thaliana* (Sugio et al., 2011). AY-WB was predicted to secrete 56 secreted proteins (also known as SAPs) as putative effectors (Bai et al., 2009). Some of these SAPs are mobile and affect host cellular processes in adjacent companion cells or distal meristem tissue (MacLean et al., 2011). So far, all characterized SAPs (SAP11, SAP54, and SAP05) have been shown to target plant transcription factors, manipulating host cells by reprograming their transcriptome (Lakhanpaul et al., 2023). This transcription reprogramming not only facilitates colonization in the plant tissue but also insect transmission (Orlovskis et al., 2023). For example, SAP11 interacts with and destabilizes class II TCP (TEOSINTE BRANCHED1/CYCLOIDEA/PROLIFERATING CELL FACTOR) host transcription factors (TFs), resulting in leaf crinkling and stem proliferation. Homologs of SAP11 from Maize bushy stunt phytoplasma (MBSP), Wheat Blue Dwarf phytoplasma (WBD), and Apple Proliferation phytoplasma (AP) - referred to as SAP11_MBSP_, SAP11_WBD_, and SAP11_AP_, respectively also interact with TCP TFs. These interactions lead to significant alterations in leaf and shoot architecture (Wang et al., 2018b; Lakhanpaul et al., 2023). SAPs can induce additional developmental anomalies such as phyllody and bolting (Orlovskis & Hogenhout, 2016, Iwabuchi et al., 2020). For example, SAP54_AY-WB_ degrades MADS-box TFs in transgenic *A. thaliana*, resulting in the formation of leafy flowers (MacLean et al., 2011).

Homologs of SAPs in JWB phytoplasma have also been studied. These effectors are named SJPs (<u>S</u>ecreted <u>J</u>WB <u>P</u>roteins). The SAP11_AY-WB_ homologs SJP1 and SJP2 promote lateral bud outgrowth by destabilizing the jujube protein ZjBRC1 and regulated auxin efflux (Zhou et al., 2021). The SAP54_AY-WB_ homolog SJP3 induces pistil reversion partly through interaction with MADS-box TF SHORT VEGETATIVE PHASE 3 (SVP3) (Deng et al., 2021; Deng et al., 2024). Despite these insights, many SJPs remain uncharacterized.

Here we identified and characterized the JWB phytoplasma effector, SJP39 (AYJ01459.1), which is the most highly expressed SJP during jujube infection. SJP39 interacts with the transcription factor ZjbHLH87 (XP_015866259.1) and stabilizes its accumulation. Overexpression of *SJP39* and *ZjbHLH87* in transgenic jujube led to stunted growth, mirroring JWB symptoms, potentially through inhibiting gibberellin (GA) signalling. This finding uncovers a molecular mechanism underlying JWB pathogenesis and opens new avenues for enhancing jujube resistance to JWB.

## Materials and Methods

### Plant materials and bacterial strains

*A. thaliana* plants were grown in a greenhouse under either long-day (16 h light/8 h dark) or short-day (10 h light/14 h dark) conditions at 22°C, with plant age calculated from the date seeds were transferred to growth chambers following stratification. *Nicotiana benthamiana* plants were grown in soil under a 16-h light/8-h dark cycle at 22°C in a growth chamber. Jujube ‘Jingzao 39’ seedlings grown in subculture media (Murashige and Skoog (MS) media supplemented with 0.3 mg L^-1^ 6-butyric acid (6-BA), 0.2 mg L^-1^ indolebutyric acid (IBA), and 0.1 mg L^-1^ gibberellic acid (GA3)) and managed under long-day (14 h/10 h (light/dark)) conditions at 25 ± 1 °C.

Diseased jujube trees (*Ziziphus jujuba* Mill.) were sampled from orchards in Changping, Beijing, and Cangzhou, Hebei. Branches and roots were collected from ‘Suanzao’ (*Z. jujuba* Mill. var. *spinosa*) and ‘Jinsixiaozao’ (*Z. jujuba* Mill. ‘Jinsixiaozao’) trees exhibiting symptoms of JWB. Three independent samples were taken from each cultivar and tissue type. *Agrobacteria*-mediated transformation in *N. benthamiana* Agroinfiltration-based transient gene expression in *N. benthamiana* leaves was performed as described with minor modifications (MacLean et al., 2014). Briefly, TFs-Myc and GFP- SJP39^32-114^ (The mature SJP39 excluding the signal peptide, 1-31 amino acids), or GFP- SJP39^32-77^ or GFP were expressed in *N. benthamiana* leaves for checking protein abundance. Proteins were separated by 12% SDS-PAGE and detected using anti-GFP (1:2000, Merck) and anti-Myc (1:2000, Merck) antibodies.

### Confocal microscopy

*SJP39* along with the *SJP39^32-77^* and *SJP39^77-114^* truncates, were cloned into the pYBA1152 vector. Constructs encoding *GFP-SJP39*, *GFP-SJP39^32-77^*, and *GFP-SJP39^78-114^*. For evaluating the impact of SJP39 on the nuclear accumulation of ZjbHLH87, the coding sequence of *ZjbHLH87* was cloned into pYBA1152 vector while mature *SJP39* and *SJP39^32-77^* were cloned into the vector pGDR-DsRed. GFP-ZjbHLH87 and H2B-CFP were co-expressed with pGDR-DsRed, DsRed-SJP39, or DsRed-SJP39^32-77^ in four replicate *N. benthamiana* plants. After 3 days post-inoculation (dpi), fluorescence was visualized using a confocal laser scanning microscope as described above. The percent gain was maintained for GFP channels 25%. Four images were taken from each sample and nuclear GFP intensity was quantified using ImageJ (Brazill et al., 2018). For each image, the nuclear CFP channel was converted to binary, and nuclei were defined using the Analyse Particle feature (>10-micron, 0-1 circularity). Nuclei were added to the ROI manager and GFP intensities were measured for each nucleus in the images. Each experiment was repeated three times independently. Plasmids used in this study are listed in Table S1

### Generation of transgenic *A. thaliana* and jujube lines

To generate transgenic plants overexpressing *SJP39* and *ZjbHLH87*, the *SJP39* gene was cloned into the pYBA1152 vector, while *ZjbHLH87* was cloned into the pCAMBIA-1300 vector. Each gene construct, along with its corresponding empty vector control, was introduced into *Agrobacterium tumefaciens* strain GV3101. All *A. thaliana* lines used were in the Columbia-0 (Col-0) background, and transgenic lines were produced using the *Agrobacterium*-mediated floral dip method as described by Clough and Bent (1998).

For overexpression of *SJP39* and *ZjbHLH87* in jujube, genes were cloned into the pYBA1152 vector and subsequently transformed into *A. tumefaciens* strain GV3101, along with GFP as a control. All transgenic jujube lines were generated using the leaf disc method (Feng et al., 2010).

### Virus-induced virulence effector (VIVE) assay

The VIVE assay was conducted as previously described (Shi et al., 2020). Briefly, Polymerase Chain Reaction (PCR) products of SJP39 and SJP37 (lacking the signal peptides) were ligated into the pGR107 vector, which is driven by the strong constitutive *Cauliflower Mosaic Virus* 35S promoter (CaMV 35S) and contains the complete potato virus X (PVX) genome. The resulting construct was introduced into GV3101 and subsequently infiltrated into the leaves of 10-day-old *N. benthamiana* plants. Viral symptoms were monitored and photographed at 14 days post-inoculation (dpi). Leaves inoculated for 2 to 6 dpi were harvested, and total protein was extracted as previously described (Li et al., 2023). Viral titer was examined by western blotting using an antibody specific to the PVX coat protein (CP).

### Yeast two-hybrid (Y2H) screening

In the Y2H screen, the *SJP39* gene, excluding its secretory signal peptide, was cloned into the pDEST32 plasmid using the Gateway system (Thermo Fisher Scientific) and screened against a 1956 *A. thaliana* transcription factor library (pDEST22-AtTF), as previously described (de Folter & Immink, 2011; Pruneda-Paz et al., 2014).

For Y2H assays, *SJP39*, along with its truncated forms *SJP39^32-77^* and *SJP39^78-114^*, were cloned into pDEST32, while the coding sequences of *AtbHLH87* and *ZjbHLH87* were cloned into pDEST22. The recombinant vectors were co-transformed into the AH109 strain, and yeast cells containing both bait and prey vectors were grown on SD-LTH medium supplemented with 10 mM 3-AT at 28°C for 3-5 days. Empty pDEST22/32 vectors and the pDEST22- AtTCP13(AT3G02150) /pDEST32-SAP11 combination were used as negative and positive controls, respectively (Sugio et al., 2011).

### In vitro pull-down assay

Mature *SJP39* and truncates *SJP39^32-77^* and *SJP39^78-114^* were cloned into the pGEX4T-1 vector containing the glutathione S-transferase (GST) tag, while *ZjbHLH87* and *AtbHLH87* were cloned into the pET-30a vector containing histidine (his) tag. These constructs, along with empty vectors, were expressed in *E. coli* BL21. Purified GST, GST-SJP39, GST- SJP39^32-77^, GST- SJP39^78-114^, His-ZjbHLH87, and His-AtbHLH87 fusion proteins were incubated with glutathione agarose (Thermo Scientific) at 4°C for 3 hours. After incubation, the beads were washed four times with wash buffer (20 mM Tris-HCl, pH 7.4, 1 mM EDTA, 200 mM NaCl, 1 mM DTT) and proteins were eluted using elution buffer (20 mM Tris-HCl, pH 7.4, 200 mM NaCl, 1 mM DTT, 10 mM glutathione). Both input and pull-down samples were separated by 12% SDS-PAGE and analysed by immunoblotting with anti-GST antibody (1:2000, TransGen Biotech) or anti-his antibody (1:2000, Thermo Scientific).

### Co-immunoprecipitation (Co-IP) assay

For Co-IP assays, the coding sequences of mature *SJP39*, *SJP39^32-77^*and *ZjbHLH87*, *AtbHLH87* were cloned into pYBA1152 vector and pCAMBIA-1300 vectors, respectively. The resulting constructs were transformed into *Agrobacterium tumefaciens* GV3101 and co-infiltrated into *N. benthamiana* leaves. After two dpi, approximately 2 g of leaf material was ground into a fine powder in liquid nitrogen and homogenized in 4 mL of extraction buffer containing 2% PVPP, 10% glycerol, 50 mM Tris (pH 7.5), 1 mM EDTA (pH 8.0), 150 mM NaCl, 0.1% NP-40, 1% (v/v) cOmplete-EDTA-free protease inhibitor cocktail (Sigma), 25 mM MG132 (Merck), 1 mM PMSF and 1 mM DTT. Total protein extracts were incubated with 10 μL of GFP-Trap magnetic agarose beads (Ychromotek, gtma-400) for 1.5 hours at 4°C. The beads were washed three times with wash buffer (same as extraction buffer, minus PVPP) and resuspended with 30 μL wash buffer and boiled for 5 minutes in10 ul 4×SDS loading buffer containing 10 mM DTT. Both input and immunoprecipitated samples were separated by 12% SDS-PAGE and detected using anti-GFP (1:2000, Merck) and anti-Myc (1:2000, Merck) antibodies.

### RNA isolation and RT-qPCR analysis

Total RNA was extracted from jujube transgenic lines using the FastPure Cell/Tissue Total RNA Isolation Kit V2 (Vazyme, Nanjing, China). For complementary DNA (cDNA) synthesis, 1 μg of the isolated RNA was utilized with the HiScript IV 1st Strand cDNA Synthesis Kit using the DNase I treatment to remove genomic DNA (Vazyme). Quantitative PCR (qPCR) was conducted with the HiScript II One Step qRT-PCR SYBR Green Kit (Vazyme), following the reaction conditions and program outlined by Zhou et al. (2021). The relative expression levels were calculated using the 2^−ΔΔCt^ method (Livak and Schmittgen, 2001), with normalization against the *ZjACT1* gene (LOC107413530) for all jujube-related samples and positive transgenic lines. A list of primers used in this study can be found in Table S2.

### Dual luciferase assay

For promoter cloning, genomic DNA from jujube was extracted using the DNeasy Plant Mini Kit (Qiagen, 69106). The 1021-bp promoter of *ZjKAO* (LOC107420337) and the 1106-bp promoter of *ZjGRP11* (LOC107418476) were amplified by PCR and cloned into the pGreen II 0800-LUC vector to drive expression of the firefly luciferase (Luc) gene as a reporter. Recombinant Luc vectors were individually transformed into GV3101 cells harbouring the pSoup vector which constitutively expresses Renilla luciferase (Ren) activity. Additional constructs, including AtbHLH87/ZjbHLH87-Myc, pYBA1152, GFP-SJP39 and GFP- SJP39^32-77^ were also prepared. GV3101 cultures were combined and adjusted to a final OD_600_ of 0.5 in infiltration buffer and co-infiltrated into *N. benthamiana* leaves. After 2 days, agroinfiltrated leaves were harvested, and luciferase activity was measured using the Dual-Glo Luciferase Assay System (Promega, E2920). Briefly, two leaf discs (4 mm in diameter) were collected in 2 ml tubes, ground to a powder in liquid nitrogen, and 100 μl PLS buffer (Promega, E1941) was added. After centrifuging at 4°C for 10 minutes, followed by another 5-minute centrifugation, 75 μl of luciferase assay reagent was mixed with an equal volume of supernatant in 96-well plate (Merk, CLS3922), and firefly luminescence was measured on a SpectraMax ID5 plate reader. Renilla luminescence was measured 10 minutes later using 75 μl of Stop & Glo reagent (Albert et al., 2021). Results were expressed as the ratio of firefly to Renilla luciferase activity (Luc/Ren). This experiment was repeated three times.

### Gibberellic treatment

Stem segments (∼1 cm in length) from jujube ‘Jingzao 39’ wildtype (WT), *GFP* transgenic plants, *SJP39* transgenic plants, and *ZjbHLH87* transgenic plants were excised, with all leaves removed. These stem segments were placed on Murashige and Skoog (MS) media supplemented with 0.3 mg L^-1^ 6-benzylaminopurine (6-BA) and 0.2 mg L^-1^ indole-3-butyric acid (IBA), with or without 2 mg L^-1^ gibberellic acid (GA3), and cultured for 4 weeks under long-day conditions (14 h light/10 h dark) at 25 ± 1°C. After 4 weeks, plant height was measured, and photographs were taken for documentation. Approximately 4-week-old *A. thaliana* plants expressing *GFP*, *SJP39*, and *SJP39^32-77^* were sprayed weekly with 100 µM GA3 for 3 weeks (Debeaujon & Koornneef, 2000). Plant height was measured in the sixth week. This experiment was repeated three times.

### Genome analysis of JWB phytoplasma to predict candidate effectors

The JWB phytoplasma genome (GenBank assembly accession GCA_003640545.1) was analysed following the method described by Bai et al. (2009) to identify candidate effectors. The presence of N-terminal signal peptides (SP) within predicted open reading frames (ORFs) was assessed using the SignalP program across three versions: SignalP v3.0 (https://services.healthtech.dtu.dk/services/SignalP-3.0/), SignalP v4.0 (http://www.cbs.dtu.dk/services/SignalP-4.0/), and SignalP v5.0 (https://services.healthtech.dtu.dk/services/SignalP-5.0/) as described by Petersen et al. (2011). ORFs predicted to encode transmembrane domains were identified using TMHMM v2.0 (http://www.cbs.dtu.dk/services/TMHMM/). ORFs containing transmembrane domains were excluded from further analysis. The presence of a nuclear localization signal (NLS) was determined using PSORT and PredictNLS (Cokol et al. 2000) programs. SJP15 and SJP43, identified as effectors by Deng et al. (2021), were included in this study. The list of candidates SJP effectors, SignalP and results are provided in Table S3.

### Phylogenetic analyses

For the phylogenetic analysis of bHLH87, the sequences from the VIIIb subfamily of the bHLH transcription factor family (Gao & Dubos, 2023) in *A. thaliana* and jujube were obtained from the Plant Transcription Factor Database (PlantTFDB, https://planttfdb.gao-lab.org/). Multiple sequence alignment was performed, and a phylogenetic tree was constructed using MEGA X (Tamuraet al., 2021). For the phylogenetic analysis of *Candidatus* Phytoplasma, the TimeTree of Life website (https://timetree.org/) was used (Kumar et al., 2017).

### RNA-seq analysis

Total RNA from infected jujube was extracted using the Omega Total RNA Extraction kit. Phytoplasma mRNA was enriched using the MICROBEnrich™ kit (Thermo Fisher Scientific) and Ribo-Zero rRNA Removal kit (Epicentre), respectively. RNA quality and concentration were assessed with a Nanodrop 2000 spectrophotometer. Paired end 150 bp sequencing was performed using the Illumina HiSeq™ 2500 platform. The phytoplasma data used for the analysis has been deposited into the NCBI database under the accession number PRJNA1158699.

Three 4-week-old transgenic lines carrying either *GFP*, *SJP39* or *ZjbHLH87*, were used for RNA-seq. RNA extraction was performed as above. RNA library construction and sequencing were conducted by NovaSeq (Beijing) using the DNBseq-T7 platform, with 150-bp paired-end reads.

The quality of the raw sequencing data was checked with FastQC v0.11.9 (Andrews, 2010). Subsequent statistical analyses were performed using R. To ensure high-quality sequences for mapping and downstream analyses, low quality reads and adapter had been trimmed by the sequencing facility prior to sequence delivery. RNA-seq reads were aligned with the indexed jujube bcultivar Dongzao genome assembly (GCF_031755915.1) and analysed it using HISAT2 v2.2.1. The number of reads mapped to each gene was counted using the feature counts function of the Rsubread package (Liao et al., 2019). The transgenic jujube data used for the analysis has been deposited into the ENA database (BioProject No. PRJEB81825). Differentially expressed genes (DEGs) were identified from gene counts for each sample using the DESeq2 package v1.30.1 from Bioconductor (Love et al. 2014). DEGs were selected based on |log_2_-fold change| > 2 that have an adjusted p value (padj) below a false discovery rate (FDR) cutoff of 0.05. Principal components analysis (PCA) of rlog transformed read counts was performed for all samples using the plotPCA function in DESeq2. The GO-term analysis was conducted using the biocfManager package, ViSEAGO v.1.4.0. The custom GO-terms for jujube were annotated using the Custom2GO and its annotation function. Enriched GO-terms were identified from DEG sets using the entire genome as the background gene set using the “create_topGO data” function. To test for significant enrichment, the classic algorithm and fisher exact test were used with a cutoff p value of 0.05. Redundant enriched GO terms were identified using the REVIGO web server using the very small parameter (Supek et al., 2011).

## Statistical analysis

Statistical analysis was performed in Prism 8.2. One-way ANOVA was used to analyse experimental data with more than 2 two experimental groups followed by Tukey’s multiple comparisons test, and two-tailed unpaired Student’s t test was used for other data analysis.

## Results

### SJP39 is a highly expressed effector of JWB Phytoplasma with plant manipulating activity

To identify potential virulence factors, we predicted secretion proteins, based on the presence of an N-terminal signal peptide and absence of transmembrane domains, from the genome of JWB phytoplasma (GenBank assembly accession GCA_003640545.1). This analysis revealed 72 SJP candidates (Table S3), including 43 that have been previously reported (Deng et al. 2021). The predicted SJPs were investigated for expression profiles in jujube by RNA-seq. Shoot and root tissues of infected ‘Suanzao’ (*Z. jujuba* Mill. var. spinosa) and ‘Jinsixiaozao’ (*Z. jujuba* Mill. ‘Jinsixiaozao’) were collected and analysed. RNA-seq results show that SJP3, SJP37, and SJP39 exhibited the highest expression levels in jujube (Fig. 1a), suggesting that they may play a role in colonizing jujube plants. SJP3 has been previously characterized to disrupt pistil development in *A. thaliana* and *N. benthamiana* (Deng et al., 2021; Deng et al., 2024). SJP39 was highly expressed in shoot tissues in ‘Suanzao’ whereas SJP37 was highly expressed in the roots.

**Figure 1.**
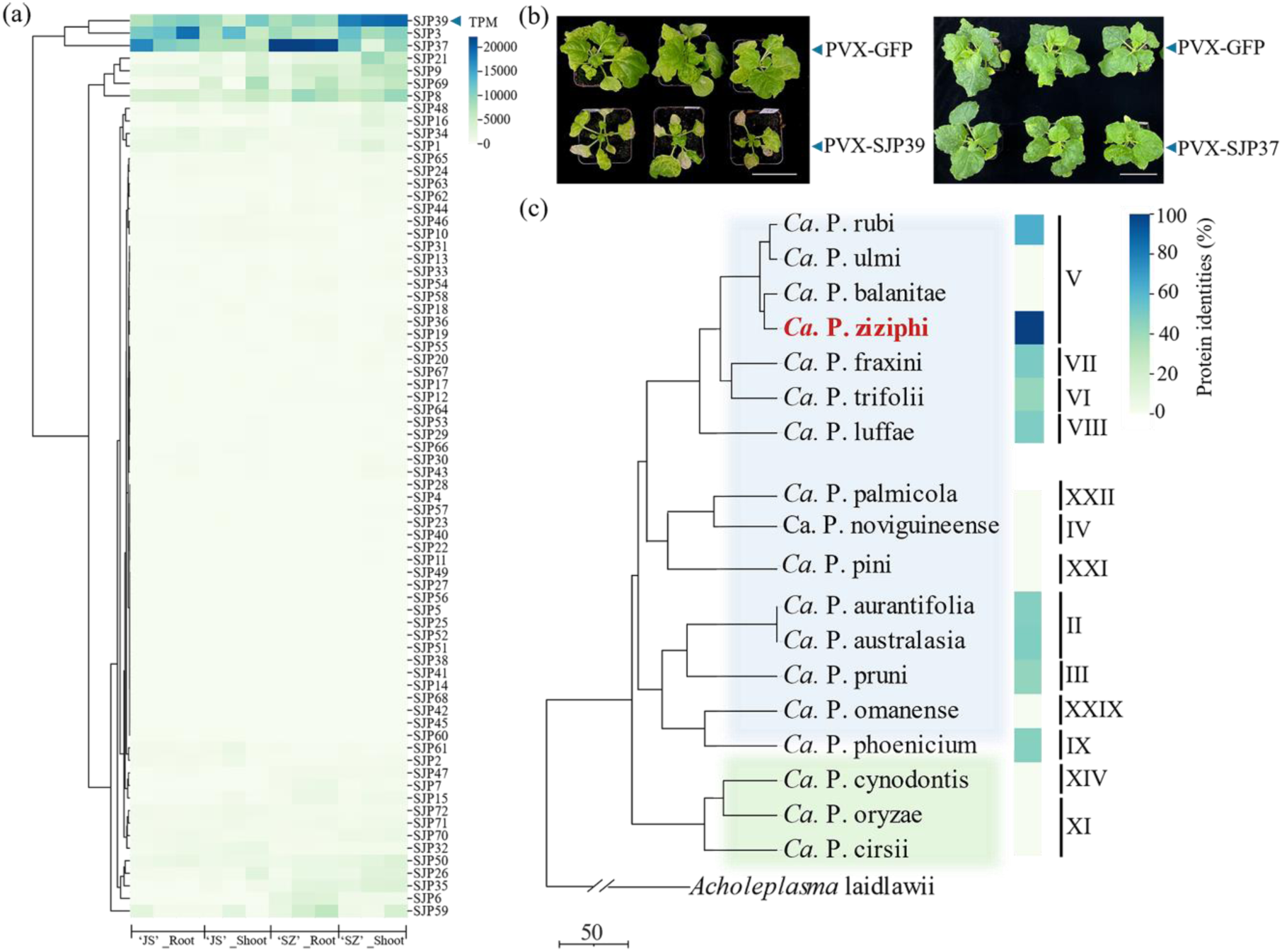
SJP39 is a putative effector produced by JWB phytoplasma. (a) SJP39 is highly expressed in JWB-infected jujube. Expression profiles of predicted Sec-dependent JWB proteins (SJPs) was determined by RNA-seq in the root and shoot tissues of two jujube (*Z. jujuba*) varieties ‘jin si xiao zao’ (‘JS’) and ‘suan zao’ (‘SZ’). (b) SJP39 enhanced necrosis after Potato Virus X (PVX) infection in *N. benthamiana*. 10-day-old seedlings of *N. benthamiana* (n = 12) were inoculated with *Agrobacterium* carrying the vectors PVX-GFP, PVX-SJP37 or PVX-SJP-39 vectors. The images were taken at 14 days post agroinfiltration. This experiment was repeated three times with similar results. Scale bar = 5 cm. (c) SJP39 homologs are produced by multiple phytoplasmas. The figure displays a time-calibrated phylogenetic tree (TimeTree) representing the evolutionary relationships among various *Candidatus* Phytoplasma (*Ca*. P.) species. Each branch corresponds to a species, and the tree scale (bottom left) indicates divergence time in millions of years (Mya), with 50 Mya as the reference. The 16S rRNA gene (16Sr) group assignments of representative phytoplasmas were determined by Davis et al. (2017). *Acholeplasma laidlawii* was used as the outgroup. The presence and absence of SJP39 homolog in each phytoplasma, as well as their amino acid sequence similarity are indicated.

We further investigated SJP37 and SJP39 for potential plant-manipulating activity in *Nicotiana benthamiana*. These two effectors were cloned into a binary Potato Virus X (PVX) vector for *Agrobacterium*-mediated transient expression. Seedling infected with PVX-SJP39 displayed chlorosis and necrosis on the inoculated leaves at 14 dpi and severe growth defects compared to plants infected with PVX-GFP or PVX-SJP37 (Fig. 1b and S1a). This was not due to enhanced viral accumulation as the titer of PVX, reflected by the level of coat protein (CP), was not increased in plant inoculated with PVX-SJP39 (Fig. S1b). In fact, the levels of PVX CP proteins were lower in these plants compared to those inoculated with PVX-GFP, indicating that SJP39 may affect plant development. To test this, we transiently expressed SJP39 in *N. benthamiana*. Interestingly, the previous phenotype did not occur in the absence of PVX (Fig. S1c, d). These results suggest that SJP39 enhances disease symptoms caused by PVX. Thus, it may function as a virulence factor and manipulate host plants.

According to the established classification scheme and a prior study on 16S rRNA gene phylogeny (Davis et al., 2017), ‘*Candidatus* Phytoplasma ziziphi (*Ca*. P. ziziphi)’ belongs to group 16SrV and is most closely related to ‘*Ca*. P. rubi,’ ‘*Ca*. P. ulmi,’ and ‘*Ca*. P. balanitae’ (Fig. 1c). To investigate the conservation of SJP39 in phytoplasmas, we searched for secreted proteins with sequence similarity across thirteen 16Sr groups known to infect diverse plant species. This analysis identified 9 SJP39 homologs among 18 *Ca.* P. species, with protein identities ranging from 41.94% to 63.16%. These findings suggest that SJP39 plays a significant role as an effector in the evolution of phytoplasmas.

### SJP39 interacts with the bHLH87 transcription factor in plants

To investigate how SJP39 manipulates plants, we determined its interacting proteins. Previous studies demonstrated that almost all phytoplasma effectors interact with host transcription factors (TFs), therefore we sought to identify plant TFs that may associate with SJP39. A yeast two hybrid screening was performed using SJP39 as the bait against a library of 1,956 TFs, which represents 78.5% of the total TFs described in *A. thaliana* (Pruneda-Paz et al., 2014) (Fig. S2a). This screen identified 22 TFs as potential SJP39 interactors (Table S4), which were further tested using pairwise assays. These experiments confirmed AtbHLH87 (AT3G21330) as the only TF that interacts with SJP39 (Fig. S2b). The potential interaction of SJP39 with a plant TF is also supported by the observation that SJP39 was predominantly located in the nuclei when expressed in *N. benthamiana* (Fig. 2a).

**Figure 2.**
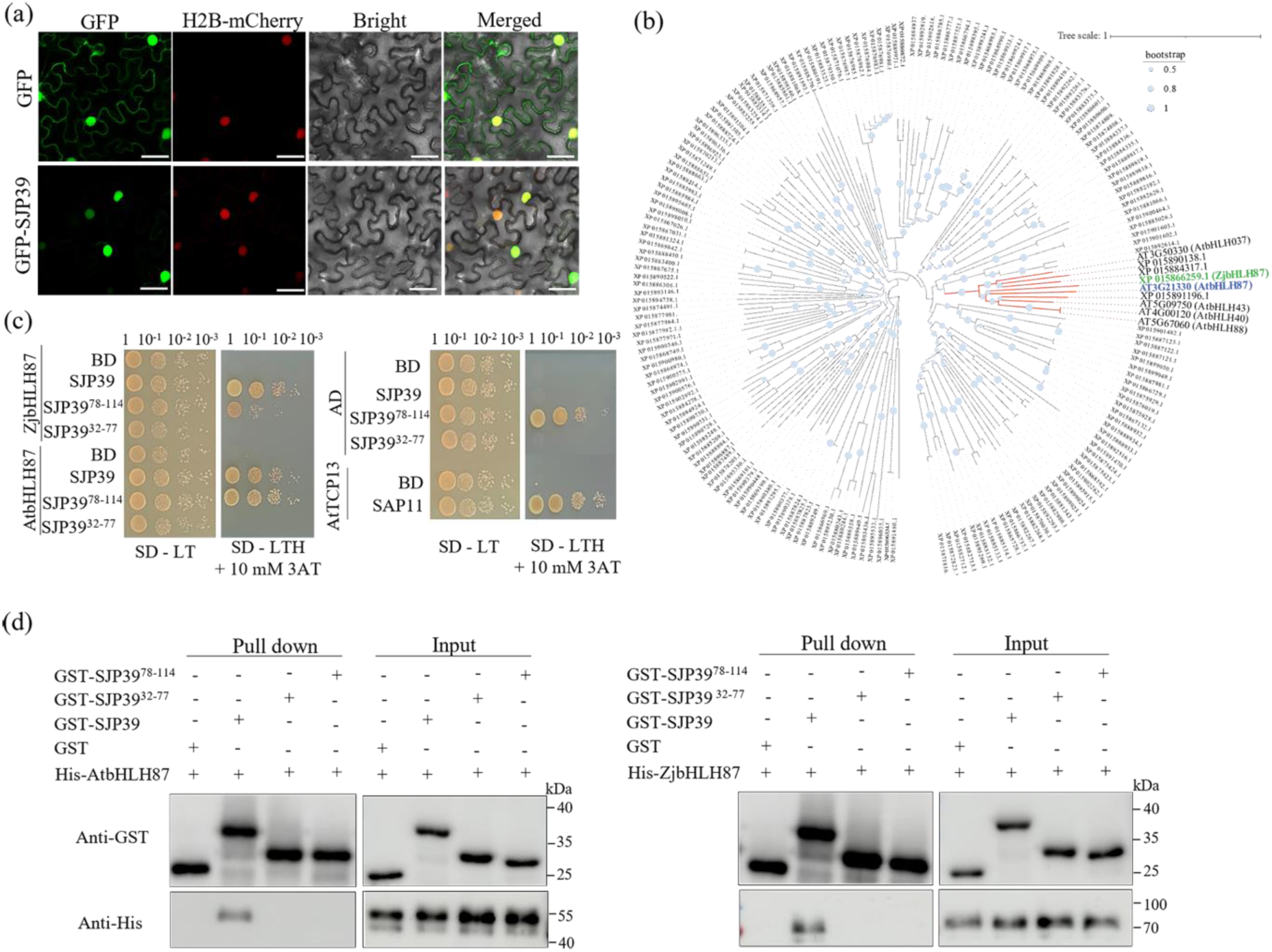
SJP39 interacts with the *A. thaliana* transcript factor bHLH87 and its jujube homolog. (a) SJP39 is located in the plant cell nuclei. Subcellular localization of GFP-SJP39 was determined in the epidermal cells of *N. benthamiana* leaves by confocal microscopy. Histone 2B (H2B)-mCherry was used as a nuclear marker. Fluorescent signals were observed at two days post Agroinfiltration. Scale bar = 50 μM. (b) Phylogenetic analysis of the VIIIb subfamily of bHLH transcription factors in *A. thaliana* and jujube. Neighbour joining method was applied to generate the phylogenetic tree using a bootstrap value of 1000. AtbHLH87 (blue), ZjbHLH87 (green) and the clade containing the bHLH87 proteins are highlighted. (c) Yeast two-hybrid (Y2H) identified bHLH87 as an interactor of SJP39. Plasmids of the bait and prey pairs were co-transformed into yeast cells and selected on double dropout (SD/-Trp/-Leu) or triple dropout (SD/-Trp/-Leu/-His) supplemented with 10 mM 3-amino-1,2,4-triazole (3-AT). Yeast co-transformed with AD-AtTCP13 and BD-SAP11 was used as a positive control. (d) SJP39 interacts with bHLH87 transcription factors in vitro. His-AtbHLH87, His-ZjbHLH87, GST-SJP39, GST-SJP39^32-77^ and GST-SJP39^78-114^ were expressed in *E. coli*. Purified protein was incubated with Glutathione Agarose. Co-precipitation of SJP39 with AtbHLH87 and ZjbHLH87 was detected by western blotting.

bHLH is a large TF family in *A. thaliana*, 171 bHLH TFs fall into 17 major groups (I-XVII) and 31 subfamilies (Heim et al., 2003). AtbHLH87 belongs to the VIIIb subfamily (Gao & Dubos, 2023). We pulled out members of the VIIIb subfamily bHLH TFs from jujube and constructed a phylogenetic tree with their *A. thaliana* homologs. This analysis revealed ZjbHLH87 (XP_015866259.1) as the closest homolog of AtbHLH87 (Fig. 2b). Y2H assays show that SJP39 also interacted with ZjbHLH87 (Fig. 2c).

SJP39 is a small protein that has 114 amino acids with the N-terminal 31 aa predicted to be the secretion signal peptide. AlphaFold2 (Jumper et al., 2021) prediction indicates two α-helices in the protein (Fig. S3). We made two truncates of SJP39, each containing one helix, and determined the region that is responsible for its interaction with bHLH87. Results from Y2H assays show that SJP39^32-77^ did not interact with AtbHLH87 or ZjbHLH87 (Fig. 2c). SJP39^78-114^ was self-active in yeast, thus we could not conclude whether this C-terminal fragment is sufficient for bHLH87 interaction.

Protein-protein interactions were further confirmed using in vitro and in planta pull-down assays. Purified proteins of His-tagged bHLH87 were incubated with GST-tagged SJP39, SJP39^32-77^, or SJP39^78-114^ and precipitated using glutathione agarose. Immunoblot analysis demonstrated that both AtbHLH87 and ZjbHLH87 could be pulled down by GST-SJP39 but not either of the truncated proteins (Fig. 2d). Co-IP assays were also conducted in *N. benthamiana* leaves co-expressing bHLH87-Myc and GFP-tagged SJP39 or SJP39^32-77^. Total proteins were immunoprecipitated using anti-GFP magnetic beads and the co-precipitation of bHLH87 proteins were detected using anti-Myc antibody (Fig. S4). Taken together, these experiments strongly suggest that the effector SJP39 directly interacts with the plant bHLH87 from both *A. thaliana* and jujube. Both the N- and C-terminal halves of SJP39 are required for this interaction.

### bHLH87 is stabilized by SJP39

Most characterized phytoplasma effectors target TFs for degradation (Sugio et al., 2011, MacLean et al., 2011, Wang et al., 2018b, Huang et al. 2021). We investigated whether SJP39 affected the stability of bHLH87 proteins. When co-expressed with GFP-SJP39 or SJP39^32-77^ in *N. benthamiana*, we observed a significantly increased accumulation of Myc-bHLH87 (Fig. 3a). In contrast, bHLH87 only accumulated to a low level when co-expressed with GFP or GFP-SJP39^32-77^. This result is also consistent with the bHLH87 input from our Co-IP assays (Fig. S4). To further validate this observation, we evaluated the abundance of bHLH87 in the plant cell nuclei using confocal microscopy. For this purpose, DsRed-SJP39 or DsRed-SJP39^32-77^ was co-expressed with GFP-ZjbHLH87 in *N. benthamiana* together with the nuclear marker H2B-CFP. In general, the nuclear localization of ZjbHLH87 remained unchanged in all samples. However, in the presence of SJP39 but not SJP39^32-77^, GFP-ZjbHLH87 exhibited significantly stronger nuclear fluorescence (Fig. 3b-c). These experiments demonstrate that SJP39 associates with plant bHLH87 in the nuclei and enhances their stability.

**Figure 3.**
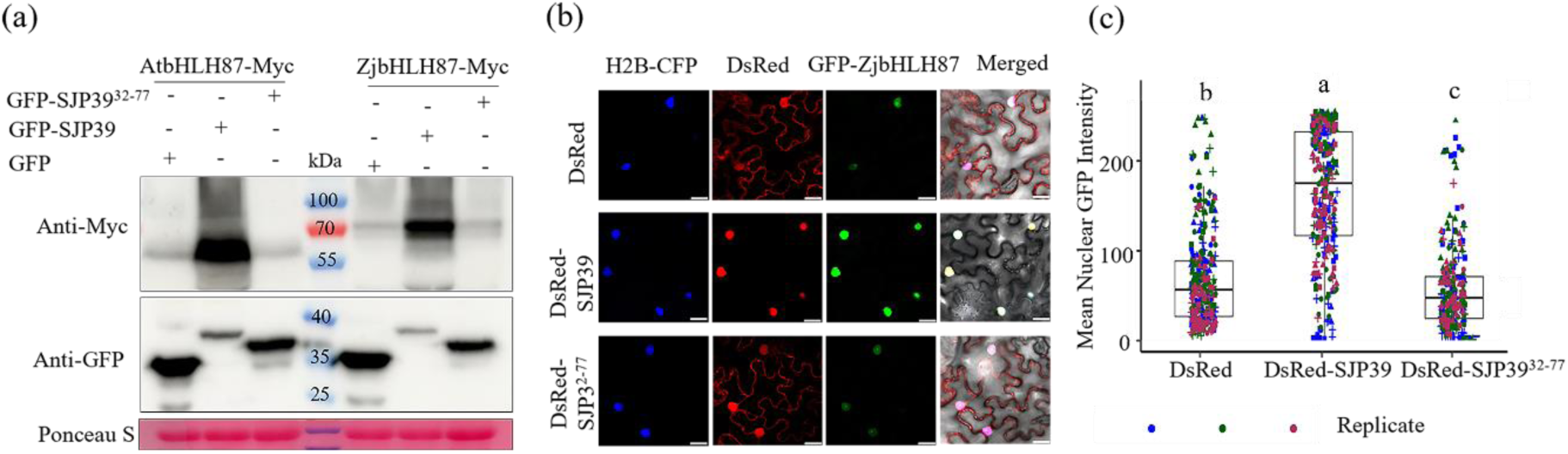
SJP39 stabilizes plant bHLH87 proteins in the nuclei. (a) SJP39 increased the accumulation of bHLH87 proteins when co-expressed in *N. benthamiana*. Myc-tagged AtbHLH87 and ZjbHLH87 were co-expressed in the presence of GFP-SJP39 or GFP-SJP39^32-77^. Protein abundance was determined by western blotting. Ponceau S staining was used to confirm equal loading. (b) SJP39 increased the accumulation of bHLH87 proteins in the nuclei. GFP-ZjbHLH87 was co-expressed with DsRed-SJP39 in *N. benthamiana*. Histone 2B (H2B)-CFP was used as a nuclear marker. The fluorescence signals were examined using a confocal laser scanning microscope after 3 days. Scale bars = 25 μm. (c) Quantification of GFP fluorescence intensity in the confocal images using Image J (n=80). Different letters indicate significant differences (one-way ANOVA, p < 0.05). Different colours represent different replicates, while different shapes correspond to distinct plants within each replicate.

### SJP39 induces developmental defects in *A. thaliana* and jujube

To further characterize the role of SJP39 and the consequence of ZjbHLH87 stabilization, we generated transgenic jujube constitutively expressing either *GFP*, *SJP39*, or *ZjbHLH87* in wild type (WT) ‘Jingzao 39’ jujube (Fig. S5a-b). Interestingly, *SJP39*-expressing seedlings exhibited smaller leaves and reduced height compared to both control lines expressing *GFP* or WT (Fig. 4a, b). Overexpressing *ZjbHLH87* also led to similar phenotypes in jujube seedlings (Fig. 4c, d). Between two independent transgenic lines, line # 4 exhibited more severe phenotypes compared to line #16, consistent with the higher expression level of *ZjbHLH87* in line #4 (Fig. S5b). JWB infection causes short internodes, yellowing, and the development of abnormally small leaves in jujube trees (Yang et al., 2023). The growth defects observed in transgenic jujube resemble symptoms observed in JWB diseased trees, suggesting that SJP39 may contribute to virulence.

**Figure 4.**
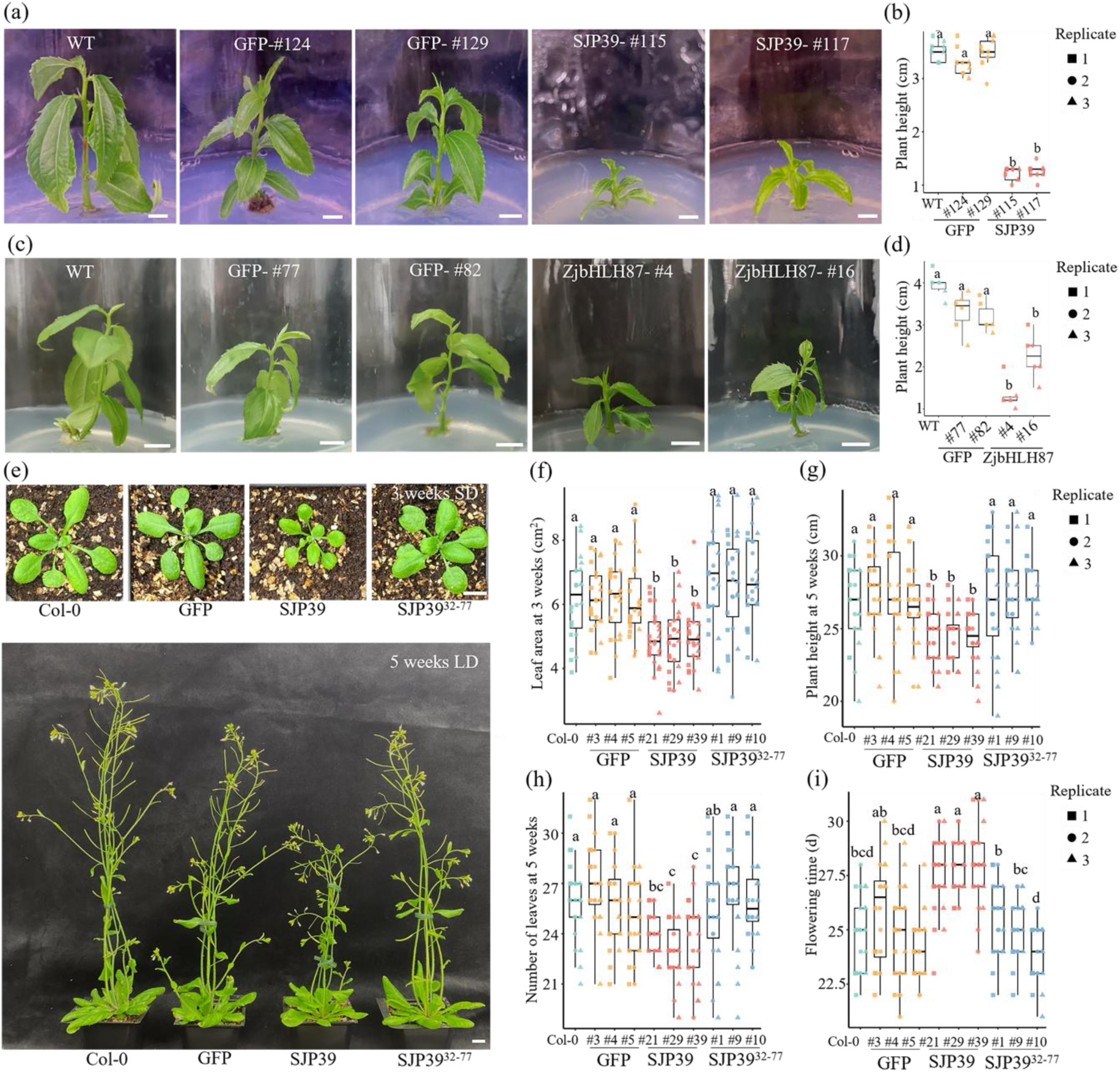
Expression of *SJP39* induces developmental defects in jujube and *A. thaliana*. (a-b) Jujube seedlings were stunted when expressing *SJP39*. 5-week-old seedling of two independent jujube lines expressing *SJP39* or *GFP* were imaged and their height quantified (n=6). Scale bar = 1 cm. WT = wildtype. Different letters indicate significant differences (one-way ANOVA analysis, p < 0.05). (c-d) Overexpressing *ZjbHLH87* also led to stunting of jujube seedlings. (e-i) Transgenic *A. thaliana* expressing *SJP39* exhibited developmental defects. (e) Representative images of transgenic *A. thaliana* expressing *SJP39* or *GFP* at 3 weeks (top) or 5 weeks (bottom) after germination. Scale bars = 1 cm. Rosette leaf area of 3-week-old plants (f), plant height (g), leaf numbers (h) of 5-week-old plants, and flower time (i) were quantified (n=8). Different letters label significant differences (one-way ANOVA analysis, p < 0.05)

We also generated transgenic *A. thaliana* lines expressing *GFP*, *SJP39* and *SJP39^32-77^* (Fig. S5c). Similar to what was observed in jujube, *SJP39* transgenic lines have smaller rosette leaves (Fig. 4e, f). Additionally, 5-week-old plants showed significantly reduced height and leaf number compared to Col-0 or transgenic plants expressing *GFP* or *SJP39^32-77^* (Fig. 4g, h). *SJP39* expression also led to delayed flowering (Fig. 4i). These results suggest that SJP39 impairs plant growth in both jujube and *A. thaliana*, mirroring disease symptoms induced by phytoplasma infection. Over-expression of *ZjbHLH87* in jujube led to similar growth inhibition, indicating that the virulence function of SJP39 is potentially mediated by stabilizing ZjbHLH87.

### SJP39 disrupt GA signalling, potentially through interaction with ZjbHLH87

SJP39-mediated ZjbHLH87 stabilization may alter the expression of genes within its regulon. We therefore analysed genes regulated by SJP39 and ZjbHLH87 in jujube seedlings expressing *SJP39* or *ZjbHLH87* using RNA-Seq. Pairwise comparisons were made between transcriptomes of *SJP39*- or *ZjbHLH87*-expressing plants and seedlings transformed with *GFP*. Differentially expressed genes (DEGs) were defined by |log_2_ fold change (log_2_FC) | > 2 and an adjusted p-value < 0.05 (Table S5). *SJP39*- and *ZjbHLH87*-expressing plants had 378 and 871 DEGs when compared to control samples, respectively. Importantly, 81 DEGs were shared between SJP39 and ZjbHLH87, indicating that they could trigger similar alteration in gene expression (Fig. 5a). GO analysis of these shared DEGs showed enriched cellular processes associated with anatomical structure (phloem) development, developmental process, diterpenoid metabolism and lipid metabolism, which were downregulated, while cytokinin signalling was upregulated (Fig. S6, Table S6).

**Figure 5.**
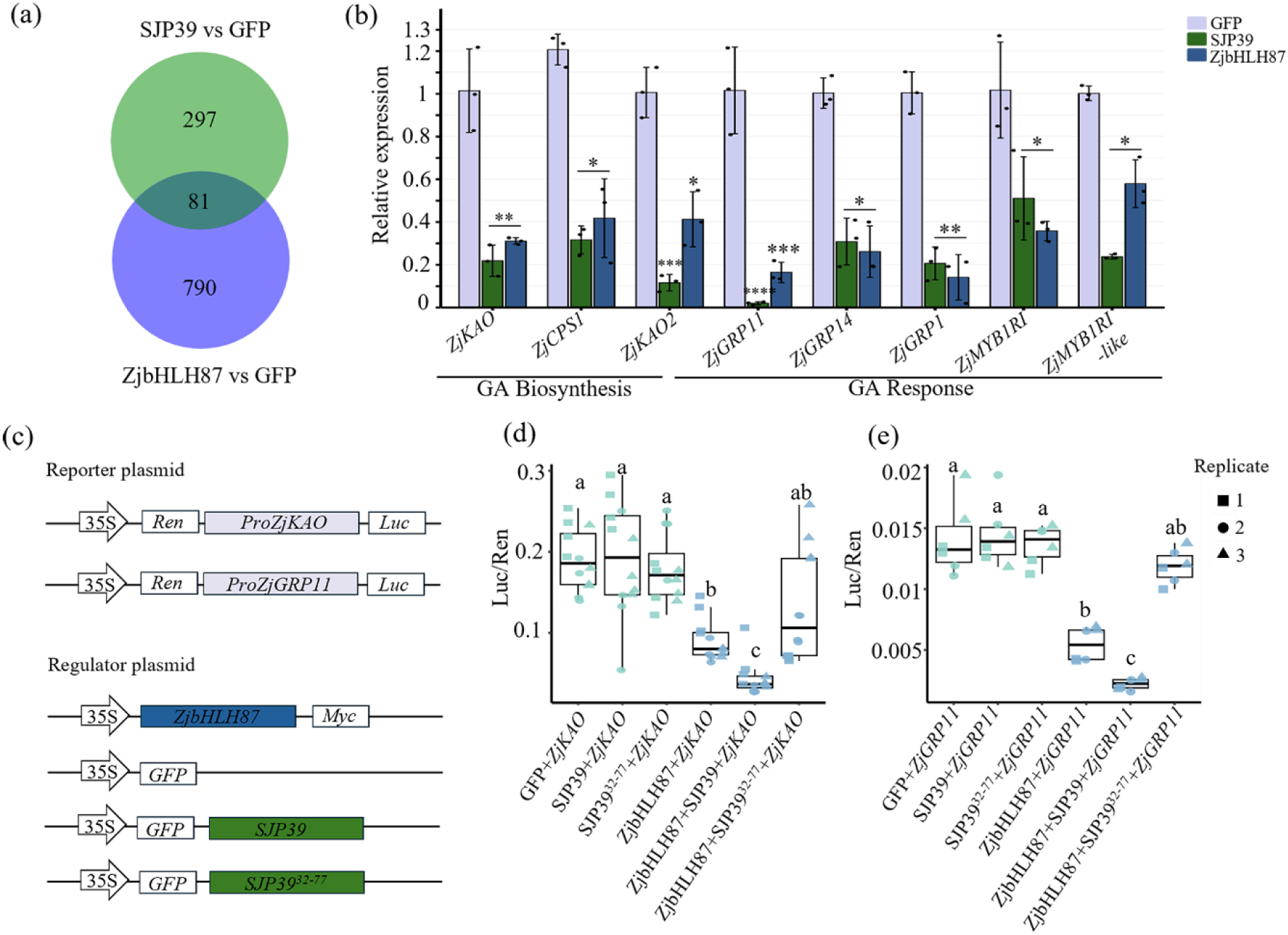
SJP39 affects GA signalling in jujube, potentially by interacting with ZjbHLH87. (a)Venn diagram of shared DEGs in jujube transgenic lines expressing *SJP39* or over-expressing *ZjbHLH87*. RNA-seq data were analyzed using pairwise comparisons between SJP39 or ZjbHLH87 and GFP. DEGs were identified as genes with |log_2_ fold change (log_2_FC) > 2 and an adjusted p-value < 0.05. A full DEG list is in Table S6. (b) Gibberellin (GA)-related genes were down-regulated in jujube seedling expressing *SJP39* or overexpressing *ZjbHLH87*. Relative expression of GA biosynthetic and responsive genes was determined using RT-qPCR (n=3). *ZjACT1* was used as an internal control. Asterisks indicate significantly different expression *p < 0.05; **p < 0.01; ***p < 0.001; ****p < 0.001 (Student’s t-test). (c-e) SJP39 enhanced the regulation of GA-related genes by ZjbHLH87. Constructs used in the dual luciferase (LUC) reporter assay (c). The *CaMV*35S promoter was used to drive the expression of Renilla luciferase (REN); while the luciferase (LUC) was cloned after the promoter of *ZjKAO* or *ZjGRP11*. The constructs were introduced into Agrobacterium, which were used for transient expression in *N. benthamiana*. LUC/REN ratios were used to indicate regulation of the *ZjKAO* promoter (d) or *ZjGRP11* promoter (e) by ZjbHLH87 in the presence of absence of SJP39. Different letters label significant differences (1-way ANOVA analysis, p < 0.05).

We were intrigued by the downregulation of the diterpenoid metabolic process in *SJP39*-and *ZjbHLH87*-expressing plants. Most DEGs within the diterpenoid metabolic pathway were downregulated (Table S6, 7). Interestingly, AtbHLH87 was reported to interact with the DELLA protein GIBBERELLIN INSENSITIVE (GAI), which is a negative regulator of gibberellin (GA) signalling (Marín-de la Rosa et al., 2014). GA is a class of diterpenoid plant hormones that is regulated by the diterpenoid pathway. Therefore, we further examined GA-related genes and examined their expression levels in *SJP39*- and *ZjbHLH87*-expressing plants using RT-qPCR (Fig. 5b). We observed significant reduction in the transcript levels of multiple genes with known functions in GA biosynthesis as well as GA responsive genes. These observations underscore a relationship between SJP39/ZjbHLH87 and GA-related pathways.

We further examined direct regulation of ZjbHLH87 on the GA-related gene expression as well as the impact of SJP39 on the regulatory activity of ZjbHLH87 using dual-luciferase (LUC) assays. Co-expression of ZjbHLH87 significantly decreased the luciferase reporter gene expression driven by the promoter of *ZjKAO* or *ZjGRP11* in *N. benthamiana* (Fig. 5c-e, S7). Furthermore, co-expression with SJP39 resulted in a further decrease in ZjbHLH87-mediated transcription suppression, while this effect was not observed in the presence of SJP39^32-77^ (Fig. 5d, e). These findings indicate that ZjbHLH87 negatively regulates the expression of *ZjKAO* and *ZjGRP11* which is enhanced by SJP39 stabilization.

To further investigate the relationship between SJP39, ZjbHLH87, and GA, we subjected *SJP39* and *ZjbHLH87* transgenic jujube lines to GA treatment. GA application significantly induced growth in WT and *GFP* lines (Fig. 6a-d). GA application was also performed on *GFP*, *SJP39*, and *SJP39^32-77^* transgenic *A. thaliana* lines (Fig. 6e). Consistent with observations in jujube, *SJP39* transgenic lines showed impaired GA responses compared Col-0 or transgenic plants expressing *GFP* or *SJP39^32-77^* (Fig. 4e, f). These results provide further evidence that the expression of *SJP39* or overexpression of *ZjbHLH87* may lead to developmental defects associated with Phytoplasma infection in jujube by inhibiting gene expression in the GA pathway.

**Figure 6.**
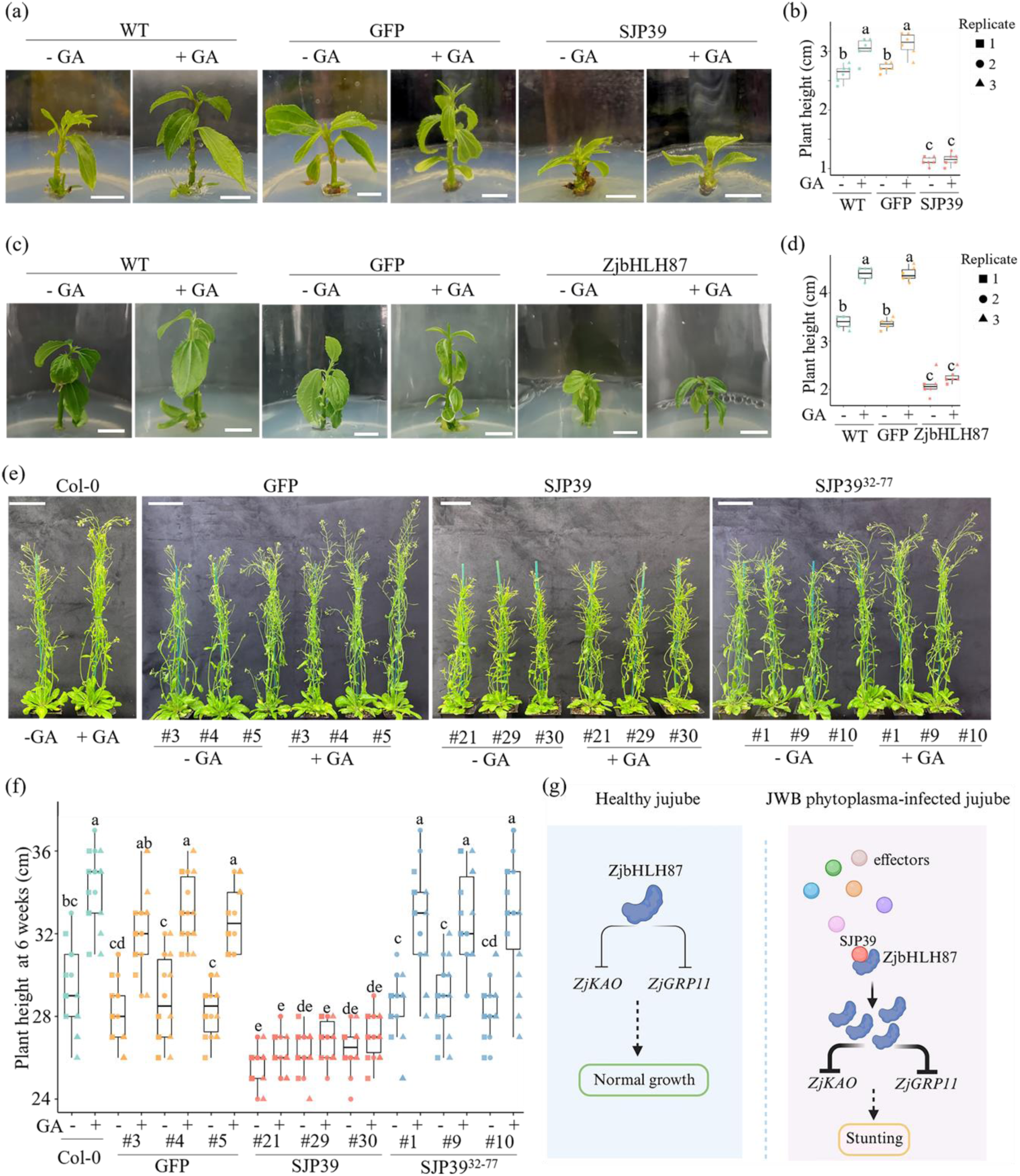
Jujube and *A. thaliana* expressing *SJP39* are insensitive to GA treatment. (a, c) Images of wild-type (WT) 5-week-old jujube seedlings or jujube expressing *GFP*, *SJP39*, or *ZjbHLH87* with (+) or without (-) 2 mg L^-1^ GA treatment. Scale bar = 1 cm. (b, d) Height of the jujube seedlings (n = 6). (e) Images 6-week-old of Col-0, or *A. thaliana* expressing *GFP*, *SJP39* or*SJP39^32-77^* with (+) or without (-) 100 µM GA treatment. Scale bar = 5 cm. (f) Height of 6-week *A. thaliana* plants. Significant differences are indicated by different letters (one-way ANOVA, p < 0.05). (g) Proposed model illustrating the virulence function of SJP39 in affecting the growth of jujube. In jujube plants infected with JWB phytoplasma, SJP39 interacts with and stabilizes the host transcription factor ZjbHLH87 in the nuclei. ZjbHLH87 acts as a negative regulator of the expression of GA-related genes such as *ZjKAO* and *ZjGRP11*. SJP39- mediated stabilization of ZjbHLH87 disrupts normal development and growth, mimicking disease symptoms observed in JWB-infected jujube. Figure created using BioRender.com.

## Discussion

Jujube is a valuable perennial fruit crop belonging to the Rhamnaceae family, with over 900 known cultivars (Zhao et al., 2019). Many of these cultivars are susceptible to JWB, a severe and often fatal disease (Liu & Zhao, 2009). Although jujube germplasm with various levels of tolerance/resistance to JWB has been identified (Song et al., 2019), a deeper understanding of the molecular interactions between phytoplasma and jujube plants that are associated with pathogenesis is required for effective disease management.

In our investigation of JWB disease, we first conducted a genome-wide effector analysis, revealing 72 candidates. Expression profiles of these candidates guided us to focus on one effector named SJP39, which is one the most highly expressed effectors in JWB phytoplasma-infected jujube trees, especially in the shoots. In addition to the JWB phytoplasma, SJP39 homologs are also produced by several other phytoplasmas that cause disease on a broad range of plants. These findings indicate a potentially important role of SJP39 in manipulating the plant hosts during phytoplasma infection.

Phytoplasma effectors predominantly target host transcription factors (TFs); and in most cases, degrade these TFs to interrupt their regulatory functions (Sugio et al. 2011, MacLean et al. 2011, Huang et al. 2021, Chen et al., 2022). SJP39 is nearly exclusively located in the nuclei when expressed in *N. benthamiana*. We identified the bHLH family transcription factor, bHLH87, as a target of SJP39. Interestingly, SJP39 stabilizes bHLH87, instead of leading to its degradation as in the case of other phytoplasma effectors to their host targets. The detailed mechanism underlying SJP39-mediated bHLH87 accumulation is currently unclear. One possibility is that SJP39 may stabilize bHLH87 by preventing its degradation through 26S proteasomes or other pathways. Alternatively, SJP39 might interact with bHLH87 to protect it from proteasomal degradation, possibly through chaperone-like functions or by modulating post-translational modifications that reduce its degradation.

bHLH TFs have been found to be targeted by effectors of other pathogens (Schwartz et al., 2017, Turnbull et al. 2017). The bacterial pathogen *Xanthomonas gardneri* produces an effector AvrHah1, which activates bHLH3 and bHLH6 transcription factors in tomato plants. These bHLH TFs regulate genes involved in pectin modification, enhancing water uptake and tissue damage, thus promoting bacterial spread (Schwartz et al., 2017). The oomycete pathogen *Phytophthora infestans* produces an effector AVR2, which induces the expression of brassinosteroid (BR)-responsive genes in potato, including the bHLH TF StCHL1. Silencing *StCHL1* reduced pathogen colonization, indicating that this TF is a positive regulator of plant immunity (Turnbull et al. 2017). Therefore, bHLH TFs with diverse functions are exploited by different pathogens to facilitate their infection.

Typical symptoms caused by phytoplasma infection include severe developmental abnormalities, such as increased lateral branching and leafy flowers. The JWB phytoplasma effectors, SJP1, SJP2 and SJP3, induced phenotypes related to these symptoms when expressed in *N. benthamiana* (Zhou et al., 2021; Deng et al., 2024). In addition, infected jujube trees often exhibit stunted growth. Interestingly, jujube seedlings expressing *SJP39* were significantly smaller than controls. Similarly, over-expression of *ZjbHLH87* also led to stunted growth in jujube, consistent with the possibility that the growth defects caused by SJP39 can be attributed to its stabilization of ZjbHLH87. While we do not have data on transgenic jujube, transgenic *A. thaliana* expressing SJP39 also showed delayed flowering, which is another symptom observed in phytoplasma-infected trees. As such, SJP39 may contribute to the disease symptom development after phytoplasma infection of jujube trees.

Transcriptome analysis of jujube expressing *SJP39* or overexpressing *ZjbHLH87* indicates a potential disruption in the gibberellins (GAs) pathway. Phytoplasma effectors are known to interfere with phytohormones and cause developmental deficiency. The effector TENGU from *Ca.* P. asteris induces stunting and floral developmental defects in *N. benthamiana* by suppressing auxin-related pathways (Hoshi et al., 2009). GA are crucial regulators of plant growth and development, particularly in controlling plant height through the promotion of stem elongation (Sun, 2010, Davière & Achard, 2013). Disruption in GA biosynthesis or signalling often leads to dwarf phenotypes, as seen in various crop species and model organisms. For example, in rice (*Oryza sativa*), mutations in the GA biosynthesis gene *GA20ox2* resulted in the semi-dwarf varieties of the “green revolution,” which exhibit reduced height (Nawaz et al., 2020). Similarly, in *A. thaliana*, mutations in *GA3ox* genes, encoding key enzymes in bioactive GA synthesis, lead to significant reductions in plant height (Hedden & Thomas, 2012; Sun et al., 2020). A suppression of GA pathway by SJP39 is therefore consistent with the stunted growth observed in the transgenic plants. A panel of GA biosynthetic and signalling genes in jujube was found to be repressed in transgenic plants expressing *SJP39*. ZjbHLH87 also negatively regulates the promoter activity of *ZjKAO* and *ZjGRP11* when co-expressed in *N. benthamiana*. Importantly, this suppression was enhanced in the presence of SJP39, likely through stabilizing ZjbHLH87. Therefore, SJP39-induced suppression of GA signalling is likely mediated by ZjbHLH87, which explains the defective growth phenotype in jujube. This is also consistent with the observation that transgenic jujube plants expressing *SJP39* or overexpressing *ZjbHLH87* were largely unresponsive to exogenous GA treatment.

Overall, our study established a role of an important JWB phytoplasma effector, SJP39, which interacts with ZjbHLH87 to enhance its negative regulation of GA signalling in jujube. This mis-regulation of the ZjbHLH87 regulon results in developmental abnormalities especially stunted growth and potentially delayed flowering, which are relevant to the disease symptoms of JWB. This finding significantly advances our understanding of the molecular mechanisms by which JWB phytoplasma causes disease on jujube and can be leveraged to develop management strategies.

## Acknowledgements

XP was supported by the National Key Research and Development Program of China (2022YFD2200404), Hebei Province Academy of Sciences Key Cooperative Unit, research on jujube witches’ broom disease pathogen, rhizosphere microorganisms, the interactions and pathogenic mechanisms with jujubes. We gratefully acknowledge Cathie Martin (John Innes Centre) for providing the pGreen II 0800-LUC and pSOUP plasmids. SY is supported by the China Scholarship Council (CSC). WM is supported by Gatsby Charitable Foundation, UKRI BBSRC Grants BB/W016788/1 and BBS/E/J/000PR9797. DHN is supported by NSF Early Career Award IOS 1942949

## Author contributions

WM and XP conceived the project and guided the execution of the experiments. SY and AL performed the experiments and analysed the data. YY and YG contributed to generating transgenic jujube lines. HN assisted with the VIVE assay and yeast two-hybrid screening. WC and WB supported the RNA-seq analysis. SY and AL prepared figures and tables. DHN guided the yeast two hybrid screening. WM, SY and AL wrote the manuscript with contributions from all authors.

## Data availability

The data supporting this study’s findings are available within the article and in the Supporting Information (Figs S1-S7; Tables S1-S7).

The phytoplasma RNA-seq data has been deposited into the NCBI database under the accession number PRJNA1158699.

The transgenic jujube RNA-seq data has been deposited into the ENA database (BioProject No. PRJEB81825).

## Supporting Information

**Fig. S1** Confirmation of SJP39 and SJP37 expression in *Nicotiana benthamiana*.

**Fig. S2** Yeast two-hybrid (Y2H) screening reveals that SJP39 interacts with AtbHLH87.

**Fig. S3** Protein structure prediction of SJP39, SJP3932-77 and SJP3978-114.

**Fig. S4** SJP39 interacts with bHLH87 using co-immunoprecipitation assay.

**Fig. S5** Confirmation of SJP39 and ZjbHLH87 expression in transgenic jujube and A. thaliana.

**Fig. S6** Shared significantly enriched Gene Ontology (GO) terms found in DEGs from pairwise comparisons.

**Fig. S7** Western blot analysis of expression of luciferase in *N. benthamiana*.

**Table. S1** DNA constructs used in this study.

**Table. S2** Primers used in this study.

**Table. S3** Predicted effector candidates from jujube witches’ broom phytoplasma (JWB).

**Table. S4** SJP39-associating transcription factors detected by Yeast-two-hybrid screening.

**Table. S5** Genes differentially expressed in transgenic jujube plants expressing SPJ39, ZjbHLH87 or GFP.

**Table. S6** GO terms enriched in genes differentially expressed in transgenic jujube lines expressing *SPJ39*, *ZjbHLH87* or *GFP*.

**Table. S7** Expression of GA biosynthetic and responsive genes in transgenic jujube lines expressing *SPJ39*, *ZjbHLH87* or *GFP*.

