## Supplemental Figure 1-7 for "A Witches’ Broom Phytoplasma effector induces stunting by stabilizing a bHLH transcription factor in plants"

### *New Phytologist* Supporting Information

The following Supporting Information is available for this article:

**Fig. S1** Confirmation of SJP39 and SJP37 expression in *Nicotiana benthamiana*.

**Fig. S2** Yeast two-hybrid (Y2H) screening reveals that SJP39 interacts with AtbHLH87.

**Fig. S3** Protein structure prediction of SJP39, SJP39^32-77^ and SJP39^78-114^.

**Fig. S4** SJP39 interacts with bHLH87 using co-immunoprecipitation assay.

**Fig. S5** Confirmation of SJP39 and ZjbHLH87 expression in transgenic jujube and *A. thaliana*.

**Fig. S6** Shared significantly enriched Gene Ontology (GO) terms found in DEGs from pairwise comparisons.

**Fig. S7** Western blot analysis of expression of luciferase in *N. benthamiana.*

**Table. S1** DNA constructs used in this study.

**Table. S2** Primers used in this study.

**Table. S3** Predicted effector candidates from jujube witches’ broom phytoplasma (JWB).

**Table. S4** SJP39-associating transcription factors detected by Yeast-two-hybrid screening.

**Table. S5** *Genes differentially expressed in transgenic jujube plants expressing SPJ39, ZjbHLH87 or GFP.*

**Table. S6** GO terms enriched in genes differentially expressed in transgenic jujube lines expressing *SPJ39*, *ZjbHLH87* or *GFP.*

**Table. S7** Expression of GA biosynthetic and responsive genes in transgenic jujube lines expressing *SPJ39*, *ZjbHLH87* or *GFP*.

**Fig. S1** Confirmation of SJP39 and SJP37 expression in *Nicotiana benthamiana*. **(a)** Western blot detection of GFP-3×Flag, SJP39-3×Flag and SJP37-3×Flag protein in *N. benthamiana* infiltrated leaves using anti-Flag antibody, Coomassie Brilliant Blue (CBB) staining as a loading control for analysis. **(b)** Western blot detection of PVX coat protein (CP) in *N. benthamiana* leaves infiltrated with PVX-GFP or PVX-SJP39 using anti-CP antibody two to six days post inoculation (dpi). Equal loading was confirmed by coomassie brilliant blue (CBB) staining. **(c)** Phenotype of *N. benthamiana* leaf (n=8)-5 days post agroinfiltration with expression 35S promoter-driven SJP39*.* Scale bar = 1 cm. **(d)** Western blot detection of 35S promoter-driven GFP and GFP-SJP39 protein using anti-GFP antibody, Ponceau S staining was used to confirm equal protein loading.

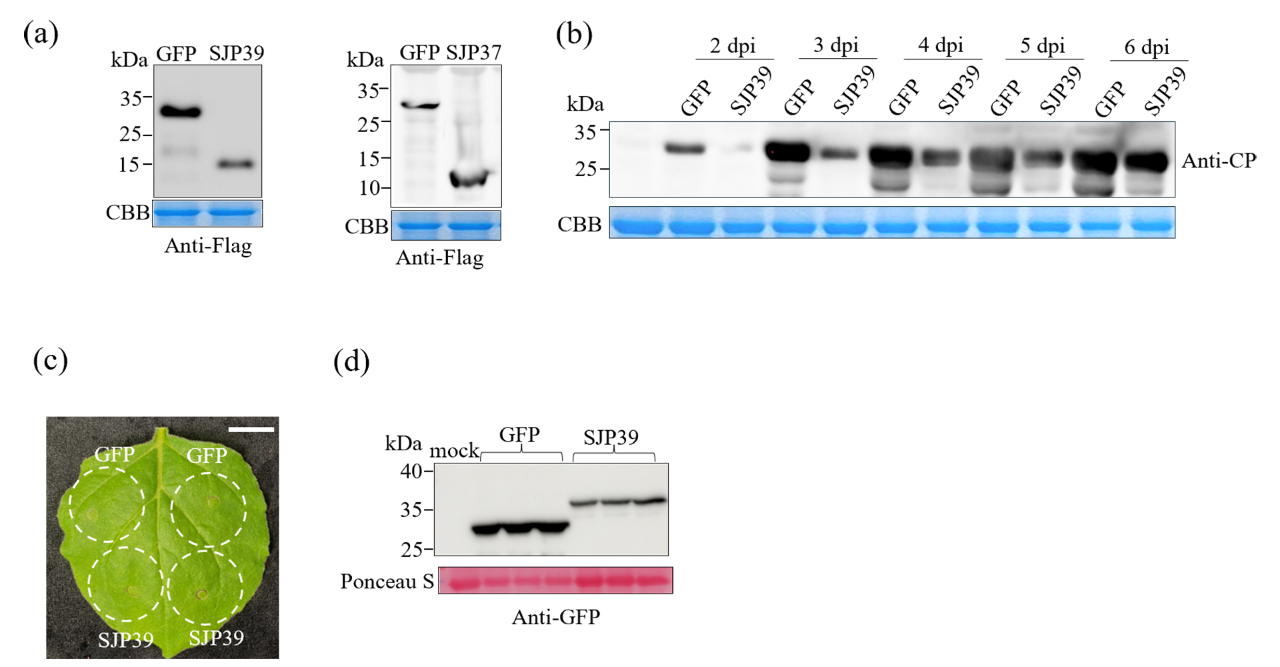

**Fig. S2** Yeast two-hybrid (Y2H) screening reveals that SJP39 interacts with AtbHLH87 **(a)** Diagram of the Y2H screening library. Blue arrows indicate the GAL4 activation domain (AD). Green arrows represent SJP39. Purple arrows denote the GAL4 DNA binding domain (BD). Pink arrows represent *A. thaliana* transcription factors. The diagram created using BioRender.com **(b)** Yeast two-hybrid (Y2H) screening identified AtbHLH87 as an interactor of SJP39. Plasmids of the bait and prey pairs were co-transformed into yeast cells and selected on double dropout (SD/-Trp/-Leu) or triple dropout (SD/-Trp/-Leu/-His) supplemented with 10 mM 3-amino-1,2,4-triazole (3-AT). Yeast co-transformed with AD-AtTCP13 and BD-SAP11 was used as a positive control.

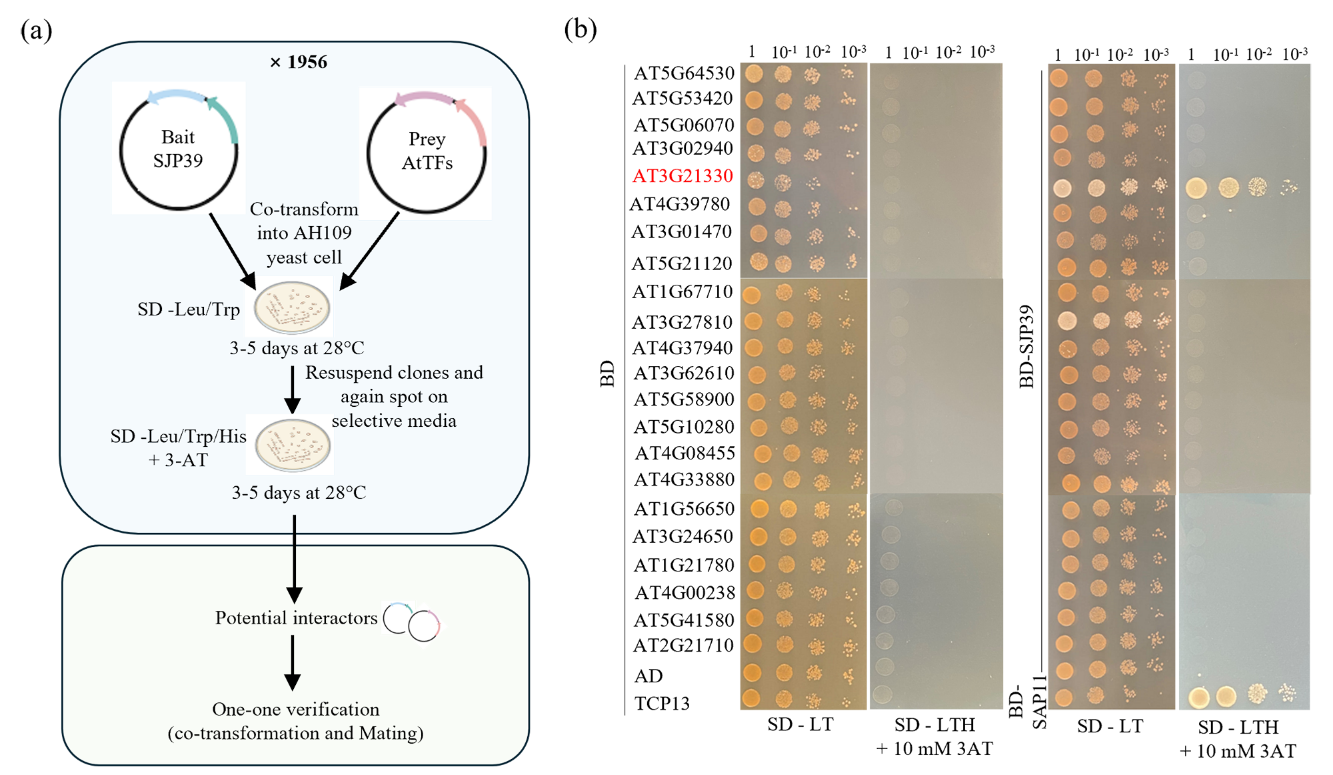

**Fig. S3** Protein structure prediction of SJP39, SJP39^32-77^ and SJP39^78-114^.

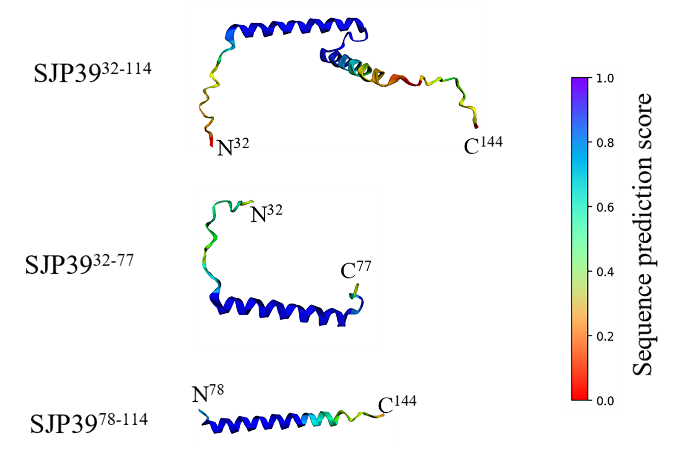

**Fig. S4** SJP39 interacts with bHLH87 using co-immunoprecipitation assay. AtbHLH87-Myc, ZjbHLH87-Myc, GFP-SJP39 and GFP-SJP39^32-77^ were expressed in *N. benthamiana*. The immune complexes were immobilized on anti-GFP magnetic beads, and the co-precipitation of SJP39 with AtbHLH87 and ZjbHLH87 was examined by western blotting using Myc antibodies. Ponceau S staining was used to confirm equal protein loading. The blue triangles indicate the correct target bands.

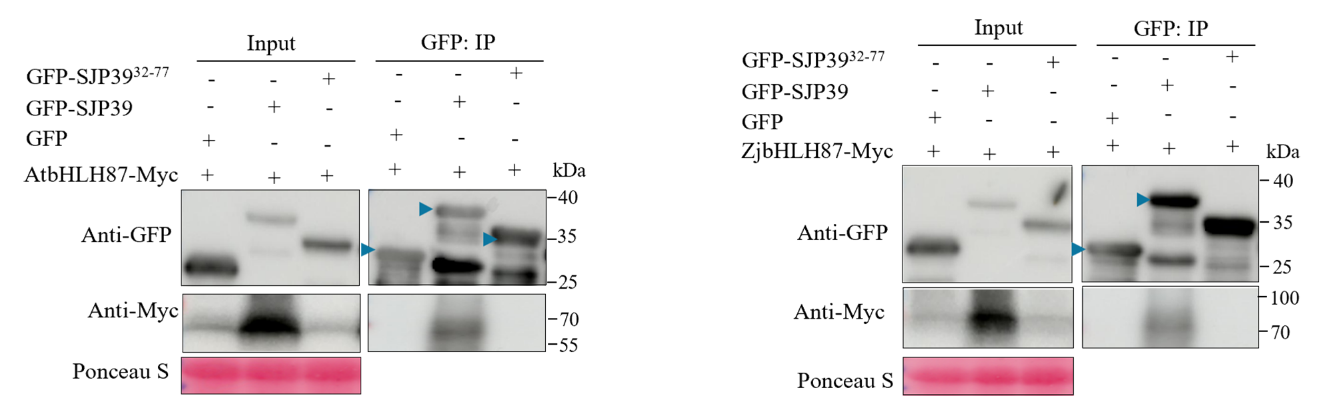

**Fig. S5** Confirmation of SJP39 and ZjbHLH87 expression in transgenic jujube and *A. thaliana*. **(a)** Western blot analysis of the GFP and SJP39 expression in transgenic jujube using anti-GFP antibody. Coomassie Brilliant Blue (CBB) staining as a loading control for analysis. The blue triangles indicate the correct target bands. **(b)** Relative expression levels of *ZjbHLH87* transgenic jujube. Transcript levels of *ZjbHLH87* measured by RT-qPCR were normalized to levels in WT control using *ZjACT1* as an endogenous control. Data are means ± SD (n = 3). Asterisks indicate significant differences (Student's t-test, *p < 0.05, **p < 0.01). (c) Western blot analysis of the expression in transgenic *A. thaliana* lines expressing *GFP*, *SJP39* and *SJP39^32-77^* using anti-GFP antibody. Ponceau S staining was used to confirm equal protein loading. The blue triangles indicate the correct target bands.

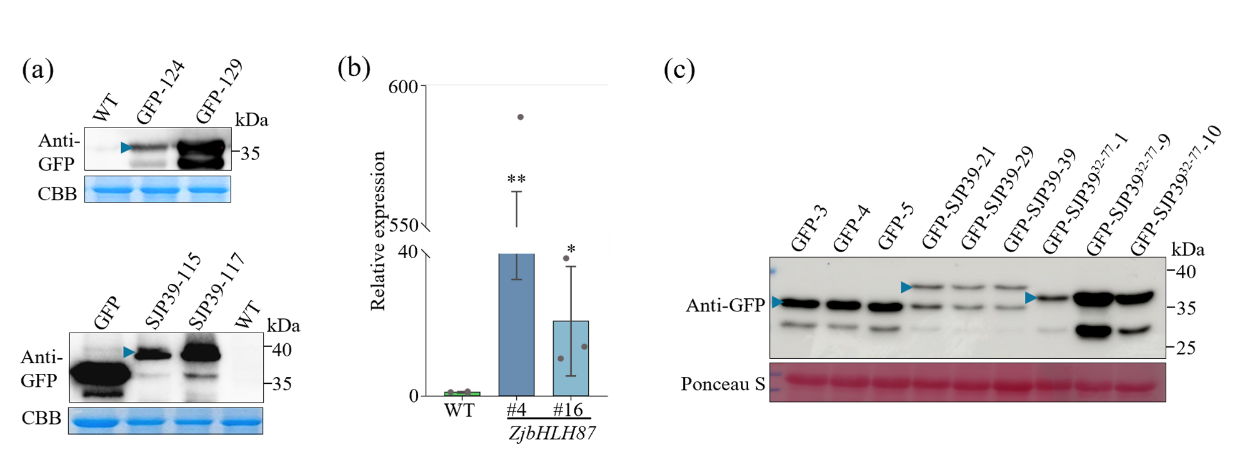

**Fig. S6** Shared significantly enriched Gene Ontology (GO) terms found in DEGs from pairwise comparisons. Significantly enriched genes for each GO terms were plotted based on their log_2_FC in SJP39 compared to GFP (green), and ZjbHLH87 compared to GFP (blue). For a full GO list, see Table S6. Asterisks indicate log_2_FC values significantly different from 0 (two-tailed t test at * p < 0.05, ** p < 0.01, *** p < 0.001).

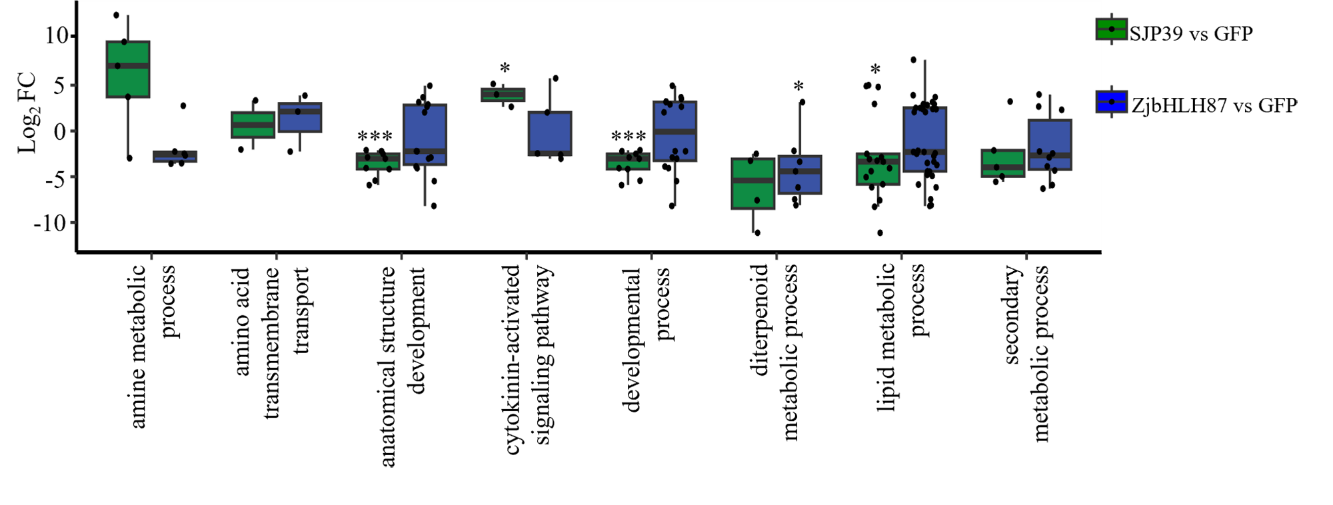

**Fig. S7** Western blot analysis of expression of luciferase in *N. benthamiana.* The constructs pGreen-luc-*proZjKAO* (left) and pGreen-luc-*proZjGRP11* (right) were co-expressed with ZjbHLH87-Myc, GFP, GFP-SJP39, or GFP-SJP39^32-77^ in *N. benthamiana* leaves. Two days post-treatment, total proteins were extracted and analysed by western blotting. Ponceau S staining was used to confirm equal protein loading.

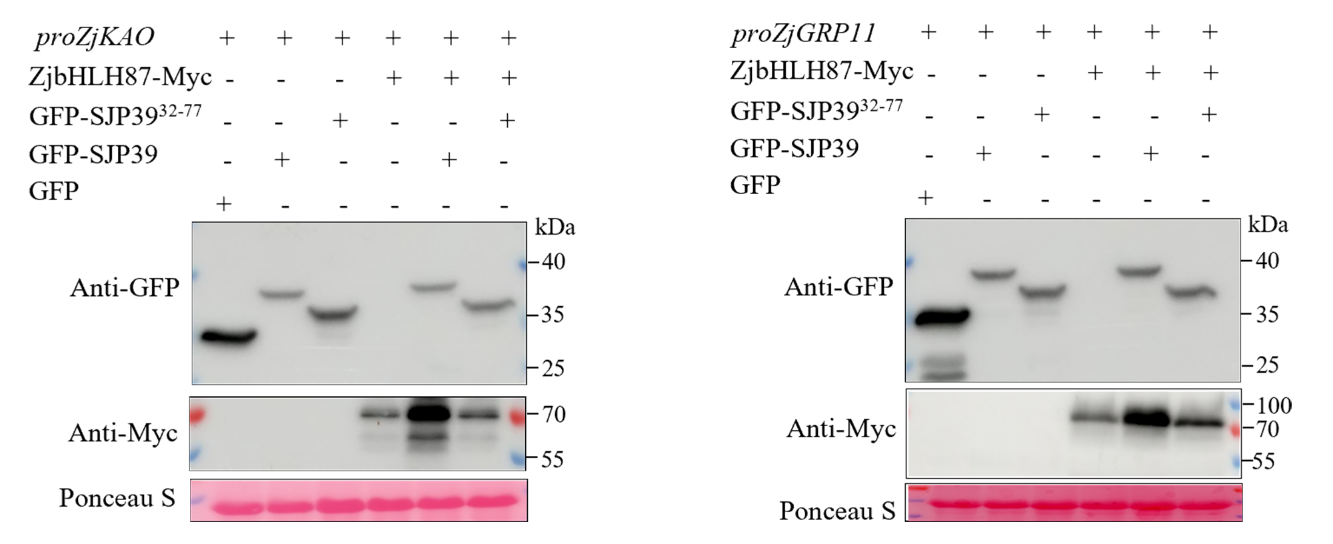
